# Allelic bias when performing in-solution enrichment of ancient human DNA

**DOI:** 10.1101/2023.07.04.547445

**Authors:** Roberta Davidson, Matthew P. Williams, Xavier Roca-Rada, Kalina Kassadjikova, Raymond Tobler, Lars Fehren-Schmitz, Bastien Llamas

## Abstract

In-solution hybridisation enrichment of genetic variation is a valuable methodology in human paleogenomics. It allows enrichment of endogenous DNA by targeting genetic markers that are comparable between sequencing libraries. Many studies have used the 1240k reagent—which enriches 1,237,207 genome-wide SNPs—since 2015, though access was restricted. In 2021, Twist Biosciences and Daicel Arbor Biosciences independently released commercial kits that enabled all researchers to perform enrichments for the same 1240k SNPs. We used the Daicel Arbor Biosciences Prime Plus kit to enrich 132 ancient samples from three continents. We identified a systematic assay bias that increases genetic similarity between enriched samples and that cannot be explained by batch effects. We present the impact of the bias on population genetics inferences (e.g., Principal Components Analysis, ƒ-statistics) and genetic relatedness (READ). We compare the Prime Plus bias to that previously reported of the legacy 1240k enrichment assay. In ƒ-statistics, we find that all Prime-Plus-generated data exhibit artefactual excess shared drift, such that within-continent relationships cannot be correctly determined. The bias is more subtle in READ, though interpretation of the results can still be misleading in specific contexts. We expect the bias may affect analyses we have not yet tested. Our observations support previously reported concerns for the integration of different data types in paleogenomics. We also caution that technological solutions to generate 1240k data necessitate a thorough validation process before their adoption in the paleogenomic community.

## INTRODUCTION

One of the major challenges faced when working with ancient DNA is the high proportion of exogenous DNA contamination present in the DNA extract. This contamination is mostly from microbes that invade the organism post-mortem or are present in the soil where the specimen was buried. It can also be introduced during laboratory experiments. To address this challenge, one method that has become popular in paleogenomic studies is the in-solution enrichment of target genomic regions using pre-designed oligonucleotides as molecular “probes” or “baits”. This technique increases the proportion of endogenous DNA of interest in a sequencing library, reducing sequencing cost and leading to larger quantities of comparable data across individual samples compared to using shotgun sequencing. Ultimately, the increase in analytical power resulting from target enrichment has enabled large-scale ancient human population genetic (i.e., paleogenomic) studies (Fumagalli, 2013).

In human paleogenomic studies, in-solution enrichment assays can be designed to target complete mitochondrial genomes (Briggs et al., 2009; Brotherton et al., 2013; Llamas et al., 2016; Maricic et al., 2010), the non-recombining portion of chromosome Y (Cruz-Dávalos et al., 2018; Rohrlach et al., 2021), genome-wide SNPs (Fu et al., 2015; Haak et al., 2015; Mathieson et al., 2015), exomes (Castellano et al., 2014), autosomes (Fu et al., 2013), or complete genomes (Ávila-Arcos et al., 2015; Carpenter et al., 2013). In 2012, Patterson and colleagues proposed a molecular bait design that used a specific ascertainment strategy to enable unbiased population genetics studies of global human populations through time (Patterson et al., 2012). The first baits using this design were applied in a handful of landmark paleogenomic studies of European populations (Fu et al., 2015; Haak et al., 2015; Mathieson et al., 2015). These bait sets were subsequently restricted to 1240k genome-wide SNPs, which paved the way for the generation of thousands of individual datasets and launched the paleogenomic era (Marciniak & Perry, 2017; Olalde & Posth, 2020; Skoglund & Mathieson, 2018). Hereafter, we will refer to this set of baits as the ‘legacy 1240k reagent’.

Since the initial publication of the molecular baits sequences in 2015, the legacy 1240k reagent was only available to a handful of laboratories through a commercial agreement. Subsequently, in 2021 two biotechnology companies—Daicel Arbor Biosciences and Twist Bioscience—released commercial in-solution enrichment kits that target the same set of 1240k SNPs (Rohland et al., 2022). In particular, the release of the suite of myBaits Expert Human Affinities kits by Daicel Arbor Biosciences aimed to make the 1240k enrichment assay accessible to all research groups worldwide, including those with limited human resources and funding. Importantly, the molecular design of both the new bait sets differs from the legacy 1240k design, with the differences specific to the Twist reagents described in a recently published study (Rohland et al., 2022). While the bait design used in the myBaits Expert Human Affinities kits is also known to differ from both the legacy 1240k and Twist assays (Daicel Arbor Biosciences, personal communication), it is a proprietary product and the specific details are unknown to the present authors. The bait design and associated laboratory protocol steps of any enrichment reagent are key to its efficacy to enrich endogenous DNA and to achieve an unbiased enrichment that respects the true proportions of reference and alternate alleles at heterozygous sites.

Here, we report genomic data from 132 ancient human individuals from western Eurasia, South and Central America (from periods preceding the colonial contact period between these populations), which were generated using myBaits Expert Human Affinities Kit “Prime Plus” (Daicel Arbor Biosciences) for various independent studies. In addition to the standard set of 1240k SNPs, the Prime Plus (PP) probes target 46,218 Y Chromosome SNPs, as well as human, Neanderthal and Denisovan whole mitochondrial genomes. Notably, when performing routine population genetic analyses on the PP-generated dataset using ƒ-statistics (Patterson et al., 2012), we observed a significant excess of shared genetic drift between any two populations enriched with the PP bait set, compared with published data generated with the legacy 1240k reagent or shotgun sequencing. These unexpected affinities could not be explained by batch effects or previously reported demographic events.

In the following sections, we outline our investigation and characterisation of the bias observed in PP-enriched ancient DNA data via principal components analyses, genetic relatedness analyses and ƒ-statistics. In total, we attempted 20 strategies to reduce or negate the observed bias, which included two wet-lab protocol adjustments and 18 bioinformatic techniques, but we were unable to correct the ƒ-statistics scores to a satisfactory standard. We conclude by recommending a series of principles for researchers to apply when co-analysing PP-enriched data with 1240k legacy and shotgun (SG) data, which mitigate the assay bias by adjusting the construction of ƒ-statistics. These guidelines may also be useful for reviewers and general readers when assessing the robustness of studies that have used data generated with the myBaits Expert Human Affinities Kit “Prime Plus”.

## MATERIALS AND METHODS

The data used in this study were initially generated for various studies intended by the authors. However, we collated all independent datasets into one overarching dataset after identifying some problems with Prime Plus data, resulting in the present study.

### Samples

We generated PP-enriched data for a total of 132 ancient samples, sourced from Spain, Mesoamerica, Peru, Bolivia and Chile. All samples from the Americas pre-date 16th-century European contact (see Table S2 for sample metadata). In our analyses, we use a comparative dataset of 115 shotgun data samples and 428 legacy 1240k samples (Bergström et al., 2020; Bongers et al., 2020; de la Fuente et al., 2018; González-Fortes et al., 2019; Günther et al., 2015; Lindo et al., 2018; Lipson et al., 2017; Mallick et al., 2016; Martiniano et al., 2017; Mathieson et al., 2015; Moreno-Mayar et al., 2018; Nakatsuka, Lazaridis, et al., 2020; Nakatsuka, Luisi, et al., 2020; Olalde et al., 2018, 2019; Popović et al., 2021; Posth et al., 2018; Prüfer et al., 2014; Raghavan et al., 2015; Salazar et al., 2023; Valdiosera et al., 2018; Villalba-Mouco et al., 2019). A complete list of the comparative samples used in this study and relevant citations are provided in Table S3.

### Data generation

#### DNA extraction

For the samples from Spain, DNA was extracted from tooth root powder using a method optimised to retrieve highly degraded ancient DNA fragments (Dabney et al., 2013) at the Autonomous University of Barcelona (UAB). For some Mesoamerican samples, the same methods were undertaken at the Australian Centre for Ancient DNA (ACAD) at the University of Adelaide. For the rest of the samples from the Americas, DNA was extracted similarly, but with a modification to the protocol that includes a bleach pre-wash step (Boessenkool et al., 2017; Dabney et al., 2013), at the University of California, Santa Cruz (UCSC) Human Paleogenomics Lab. Table S2 provides a full summary of which lab processed which samples at each step.

#### Library preparation

For the Spanish and some Mesoamerican samples, partially UDG-treated double-stranded DNA libraries (Rohland et al., 2015) were generated at ACAD. For the Bolivian, Peruvian, Chilean and the rest of the Mesoamerican samples, partially UDG-treated single-stranded DNA libraries (Kapp et al., 2021) were prepared at UCSC. Note that the difference in library structures should not impact the enrichment bias as libraries are first denatured and then hybridised to the baits.

#### Library enrichment

Before enrichment, libraries were over-amplified in order to reach the minimum required DNA input of 500 ng per sample. Library enrichment was then performed according to the suggested myBaits protocol of two enrichment rounds with hybridisation temperature at 70°C and 20 amplification cycles in the first enrichment followed by 10 cycles in the second round. Samples were grouped in pools of 2–4 libraries according to the endogenous DNA proportions obtained from screening of the initial shotgun sequences. Samples were processed at UCSC or at ACAD (see Table S2 for a breakdown of which lab processed which samples at each step).

Additionally, two experiments that examined the utility of adjustments to standard enrichment protocols were performed on a subset of libraries at UCSC (Table 1). In experiment A, a single round of hybridisation at 70°C was performed with 12 cycles of amplification. A single round of hybridisation was also used in Experiment B but at 62.5°C and with 13 cycles of amplification.

**Table 1:** Relevant modified variables for each of the laboratory experiments.

| Variable | Standard Protocol | Experiment A | Experiment B |
| --- | --- | --- | --- |
| Enrichment rounds | 2 | 1 | 1 |
| Hybridisation temperature | 70°C | 70°C | 62.5°C |
| Amplification cycles | 20 (1<sup>st</sup> round) + 10 (2<sup>nd</sup> round) | 12 | 13 |

#### Sequencing

Libraries enriched at UCSC were paired-end sequenced on a HiSeq 4000 at Fulgent Genomics (Los Angeles, USA). Paired-end sequencing using an Illumina NovaSeq 6000 (200 cycles) was performed by the Kinghorn Centre for Clinical Genomics (Sydney, Australia) for libraries enriched at ACAD (Table S2).

#### Data processing and genotype calling

Data were processed either using nf-core/eager (Fellows Yates et al., 2021), or UCSC’s in-house processing pipeline, Batpipe (https://github.com/mjobin/batpipe). The terminal 2bp at either end of each read were trimmed to remove ancient DNA damage and then mapped to the GRCh37d5 reference genome using bwa-aln with parameters −l 1024 -n 0.01 -o 2 (Oliva et al., 2021). Other mapping parameters vary slightly between different datasets used in this study, such as mapping quality threshold 20-30 and minimum read length 20-30 (see Table S2 for details of processing parameters for each population). Pseudo-haploid genotypes were called at the enrichment target sites using the random haploid mode within pileupCaller (https://github.com/stschiff/sequenceTools).

### Population genomics assessment of capture bias

#### PCA

We computed principal components analyses using both EMU (Meisner et al., 2021) and smartPCA version 18140 in EIGENSOFT 8.0.0 (Patterson et al., 2006; Price et al., 2006) with and without projection to test how the data bias affected each type of PCA.

#### *ƒ*-statistics

*O*utgroup-ƒ_3_ and ƒ_4_ statistics were computed using the ADMIXTOOLS package (Patterson et al., 2012).

#### READ

We tested the impact of the enrichment bias on genetic relatedness using READ, i.e. Relatedness Estimation for Ancient DNA (Monroy Kuhn et al., 2018), which is a widely used PMR-based estimator for genetic relatedness for ancient DNA data.

### Filters and strategies to resolve bias

In order to estimate the effect of various bioinformatic approaches to mitigate the bias observed in ƒ_4_ statistics, we calculated ƒ-statistics in the following four forms:

1) ƒ_4_(Mbuti, Spain.PP.2rnd; Target, Peru.Shotgun)
2) ƒ_4_(Mbuti, Spain.PP.2rnd; Target, Peru.1240k)
3) ƒ_4_(Mbuti, Spain.1240k; Target, Peru.Shotgun)
4) ƒ_4_(Mbuti, Spain.1240k; Target, Peru.1240k)

In each, the target population was the Peru.PP population after application of one of the 20 strategies to reduce bias detailed below. These strategies are also summarised in Table S5.

#### SNP Missingness

Amongst the set of 1240k SNPs assayed by the PP probes, each SNP was missing at least once in our dataset. Hence, we used the full data matrix with missing genotypes and reported the number of overlapping SNPs included in each ƒ-statistic comparison where relevant.

To the target population (Peru.PP), we applied two data missingness filters using the *--geno* flag in PLINK 1.9 (Chang et al., 2015)—namely, 0.0 and 0.5, which correspond to removing SNPs with >0% or >50% missingness, respectively.

#### Minor allele frequency

We tested some simple filters for minor allele frequency (i.e., 0.3, 0.1 and 0.01) using the *--maf* flag in PLINK 1.9 (Chang et al., 2015).

#### Linkage disequilibrium

We filtered for linkage disequilibrium (LD), with the intention to remove potential artefactual linkage introduced by one bait targeting multiple SNP sites. We used the PLINK 1.9 LD pruning parameter *--indep-pairwise* with a window size of 3 variants, sliding in 1 variant steps and tested correlation threshold values of *r*^2^ = 0.1, 0.3, 0.5, 0.7 and 0.9 (Chang et al., 2015). These filters resulted in datasets of 300,838, 346,428, 379,345, 388,038, and 388,678 SNPs, respectively.

Given that each *r*^2^ threshold produced approximately 300,000 SNPs, to assess if the improvement in ƒ_4_-value was driven by the reduction in bias attributable to artefactual linkage, rather than changes to other statistical properties of the data (such as decreased power due to a reduced number of SNPs), we repeated our analyses with random sub-samples of 300,000 SNPs, replicated five times, using the PLINK 1.9 flag *--thin-count (Chang et al., 2015)*.

#### Transversions

Sequencing libraries were partially treated with UDG (Kapp et al., 2021) and the terminal two nucleotides at either end of all mapped reads were trimmed when processing the data with nf-core/eager (Fellows Yates et al., 2021). Therefore, we did not expect to observe a strong impact of ancient DNA damage in the analyses. Nevertheless, we filtered out transitions to minimise the potential impact of ancient DNA damage, which is typically characterised by C-to-T substitutions at the 5’ end of sequencing reads and C-to-T or G-to-A substitutions at the 3’ end of sequencing reads when using single-stranded or double-stranded libraries, respectively (Krause et al., 2010; Meyer et al., 2012).

#### Application of Rohland et al. 2022 filter

We applied the filter suggested by Rohland and colleagues (Rohland et al., 2022) to extract SNPs that will allow co-analysis between 1240K data generated with shotgun sequencing, or the legacy 1240k, Prime Plus or Twist enrichments.

#### Read length and mapping quality threshold

We filtered out reads that were shorter than 50 bp and/or with mapping quality less than 20, as suggested by Günther & Nettelblad to reduce reference bias when mapping paleogenomic data (Günther & Nettelblad, 2019).

#### Off-target genotypes

Given the hypothesis that the bias is caused by bait-target interactions and that the enriched library efficacy is not 100%, we called off-target (non-specific) genotypes on the assumption that they would not be impacted by any bait-mediated mechanism driving the bias. The genotyping of off-target sites was performed using a standard pseudo-haploid pileupCaller call with the dbSNP set 138 (Sherry et al., 2001) after filtering out all PP target sites. Additionally, we called genotypes that were at least 100 bp away from target sites to eliminate SNPs that may have been covered by a bait but were not the target locus.

#### Majority call pseudo-haploid and random diploid genotypes

In addition to standard pseudo-haploid calls based on the random selection of a single allele at prespecified SNPs, majority allele pseudo-haploid genotypes were also called at sites with greater than 5X depth-of-coverage using the majority call mode in pileupCaller. Finally, diploid genotypes were called using the random diploid call mode in pileupCaller, where one read is chosen at random, followed by a second chosen at random without replacement, and sites covered by only one read are considered missing.

### Investigation of allele call bias

We investigated the rate of each allele call at heterozygous SNP sites for the most common substitutions: (A/C, T/G, A/G, T/C), where the inverse substitution was also included such that i.e., A/C and C/A SNPs are collated together. The relative allele call frequencies were calculated per SNP (ignoring missing SNPs) in aggregate across all population datasets for shotgun-, legacy-1240k-and PP-enriched genotypes. SNP missingness was quantified across 100 bins with equal numbers of SNPs, and the mean allele frequency per bin was plotted against SNP missingness.

## RESULTS

### Global bias assessment

To quantify the extent of the PP-associated bias in outgroup-ƒ_3_ statistics, we calculated statistics of the form ƒ_3_(Mbuti; Target, X), where Target refers to samples that were enriched using PP and originate either from ancient Spain (Spain.PP.2rnd; unpublished) or Peru (Peru.PP.2rnd; unpublished), and X denotes a set of South American and European ancient reference populations either generated from shotgun sequencing or enriched with the PP or legacy 1240k kits.

For ƒ_3_ statistics that use Spain.PP.2rnd as the Target population (Fig 1A), all non-PP European populations (shotgun or legacy 1240k data) exhibit more shared drift with the Spain.PP.2rnd population than do the South American populations of either enrichment type (Fig 1A). However, the three South American PP populations share more genetic drift with the Spain.PP.2rnd target population relative to other non-PP South American populations. Similarly, the inter-continental relationship is preserved when Peru.PP.2rnd is used as the target population, as all South American populations have substantially more shared drift with the Peru.PP.2rnd target population than with European populations, yet the intra-continental relationships appear to be violated by the biased shared drift between PP populations (Fig 1B). Further, Spain.PP.2rnd exhibits a strong attraction toward Peru.PP.2rnd samples relative to the other Spanish non-PP populations.

**Figure 1:**
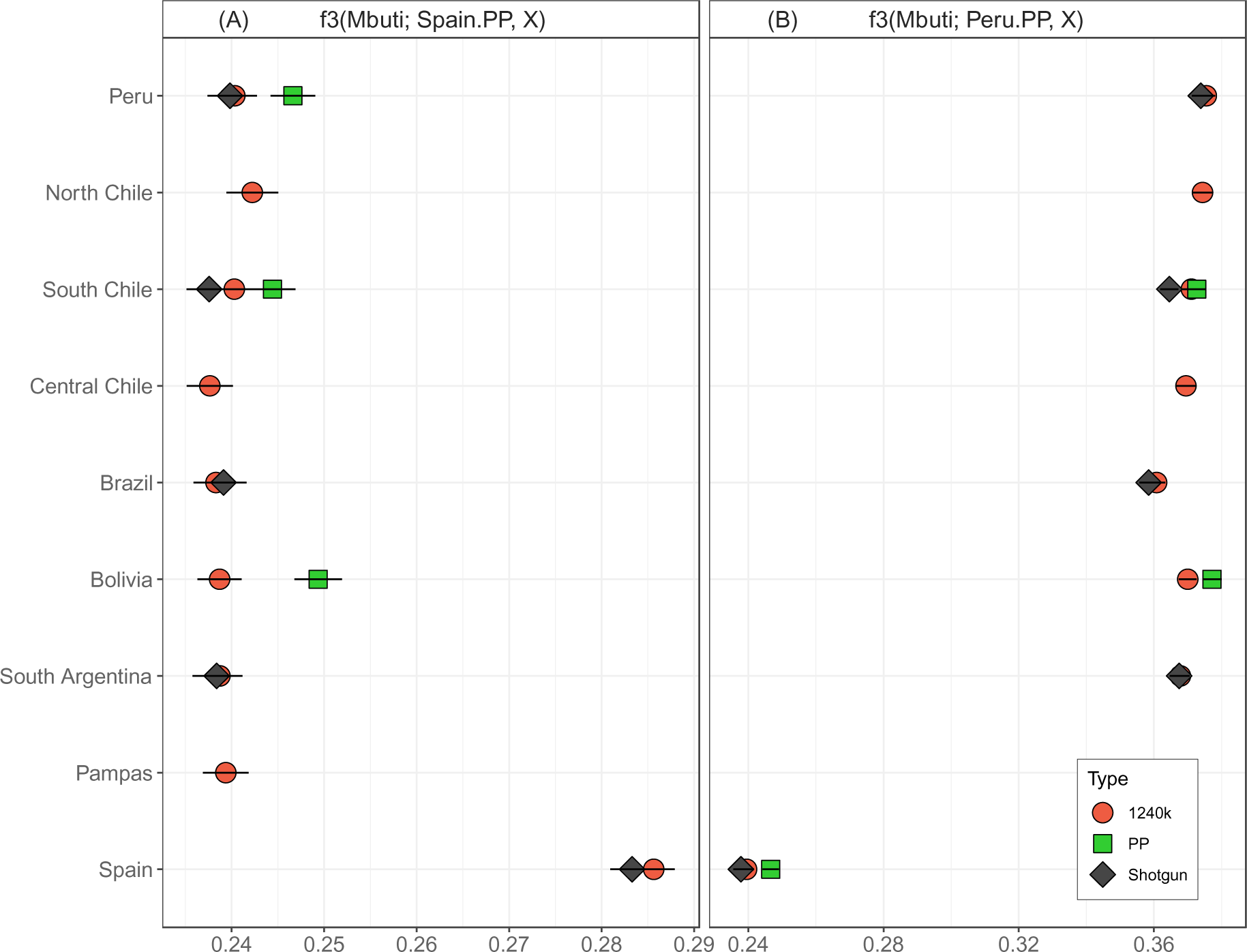
Outgroup ƒ_3_ statistics comparing ancient South American and Spanish populations by data type. Panels (A) and (B) show results for Spain.PP.2rnd and Peru.PP.2rnd as the target population, respectively. Note that not all regions have data of every type and different data types from the same region are not the same individuals but are expected to share broadly similar ancestries.

We also observe variance between 1240k and shotgun datasets, which is likely due to a combination of different ancestries and 1240k bias (Rohland et al., 2022), however in most cases the shotgun and 1240k error bars overlap, whereas the Prime Plus bias is more pronounced. Clearly, the bias observed in PP-generated data is not unique to South American populations but is ubiquitous across the highly diverged populations of Europe and South America, strongly suggesting that the bias affects all global populations.

### Principal Components Analysis

We tested the impact of PP bias on multiple PCA approaches, including the widely used smartPCA (Patterson et al., 2006; Price et al., 2006) and EMU, which is reported to handle data missingness well (Meisner et al., 2021). For each approach, we computed PCAs at continental and global scales to investigate how missingness and the PP bias manifested at different geographical scales.

#### Continental PCA

Initially, we performed a PCA using EMU on 362 ancient genome-wide datasets from South America including shotgun (n = 38), legacy 1240k (n = 223) and PP-enriched (n = 101) data (see Table S1 for a detailed list of samples). Sample clustering in the first three principal components of the PCA is driven by population structure (Fig 2A and there is no apparent bias exhibited by the data source type (Fig 2A). When we used smartPCA on the same 362 ancient genome-wide datasets (without projection onto modern diversity, due to lack of sufficient modern data), the resulting PCA produced sample clustering along PC2 that was mildly separated by the assay type for Chilean samples, though this may be driven by local ancestries (Fig 2B). Note that no individual samples were assayed with all three methods investigated, but many samples of different data types derive from the same localised ancestry groups. This result concurs with a recent study (Rohland et al., 2022), which also reported that PCA is robust to combinations of these three data types.

**Figure 2:**
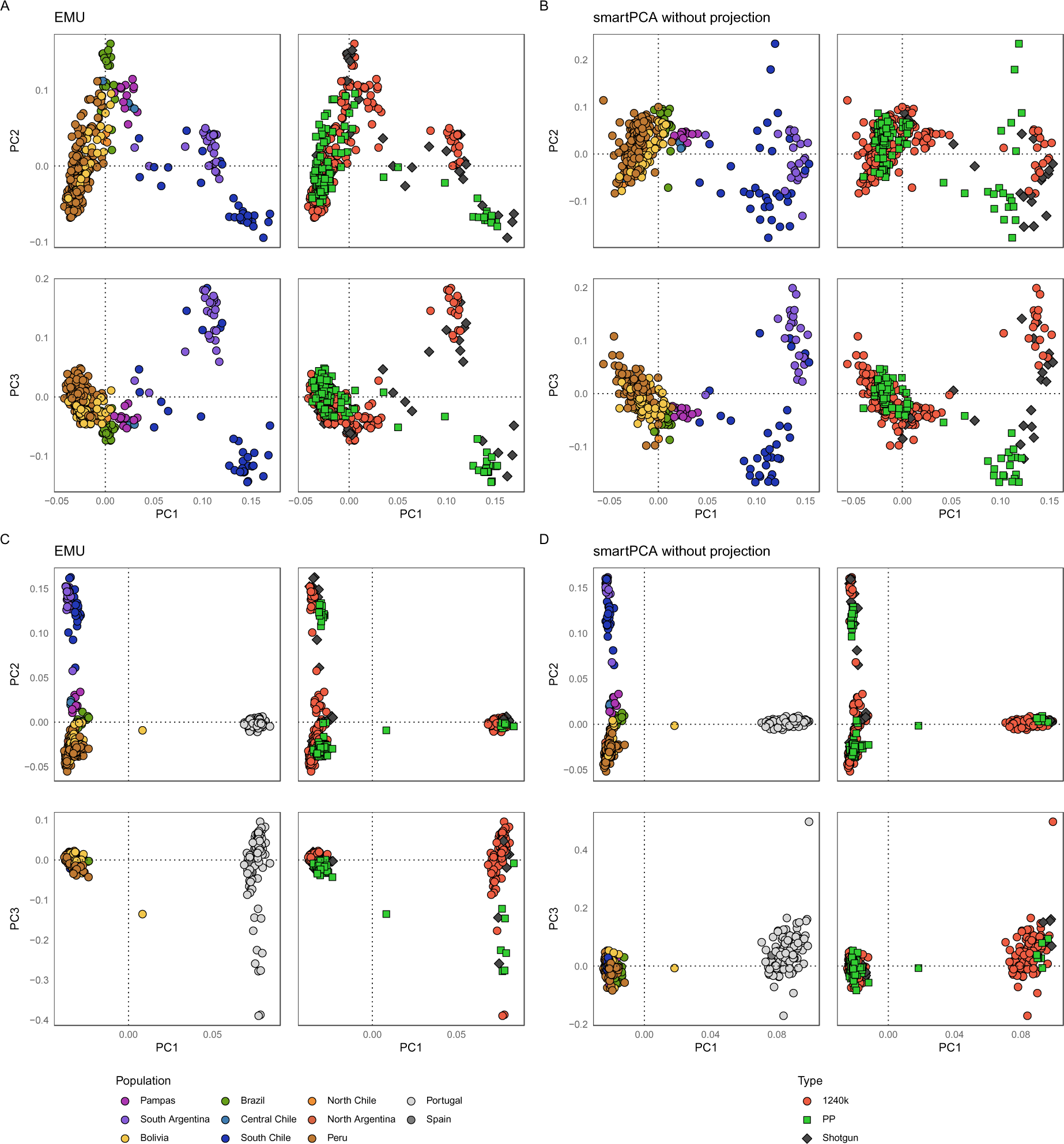
Comparing PCA methods at continental and global scales. Continental PCA (A and B) uses a collated dataset of 362 ancient genomes from South America. Global PCA (C and D) uses a collated dataset of 399 ancient genomes from South America and Iberia. Samples are coloured by region (left column of each panel) or data type (right column of each panel). (A) Continental PCA computed with EMU (Meisner et al., 2021). (B) Continental PCA computed with smartPCA, no projection (Patterson et al., 2006; Price et al., 2006). (C) Global PCA computed with EMU (Meisner et al., 2021). (D) Global PCA computed with smartPCA, no projection (Patterson et al., 2006; Price et al., 2006).

#### Global PCA

Subsequently, we performed another PCA testing both EMU and smartPCA without projection on a merged dataset of 399 ancient South American and European genome-wide datasets including shotgun (n = 38), legacy 1240k (n = 282) and PP-enriched (n = 79) data. In both the EMU and smartPCA outputs, PC1 is driven by the ancestral difference between South American and European populations, PC2 is driven by the diversity within South America and PC3 by the diversity represented in Iberia (Fig 2C and 2D).

#### Global PCA with smartPCA projection

To assess the effect of the enrichment bias on a global scale, we applied both EMU and smartPCA with projection methods to 436 ancient genome-wide datasets from South America, Mesoamerica and Europe, either generated by shotgun sequencing or enriched using the PP or legacy 1240k kits, onto a PCA of modern diversity that used 1,071 individuals from the HGDP dataset (Bergström et al., 2020) (Fig 3). Both PCAs look very similar, the ancient individuals are clustering with similar modern ancestries.

**Figure 3:**
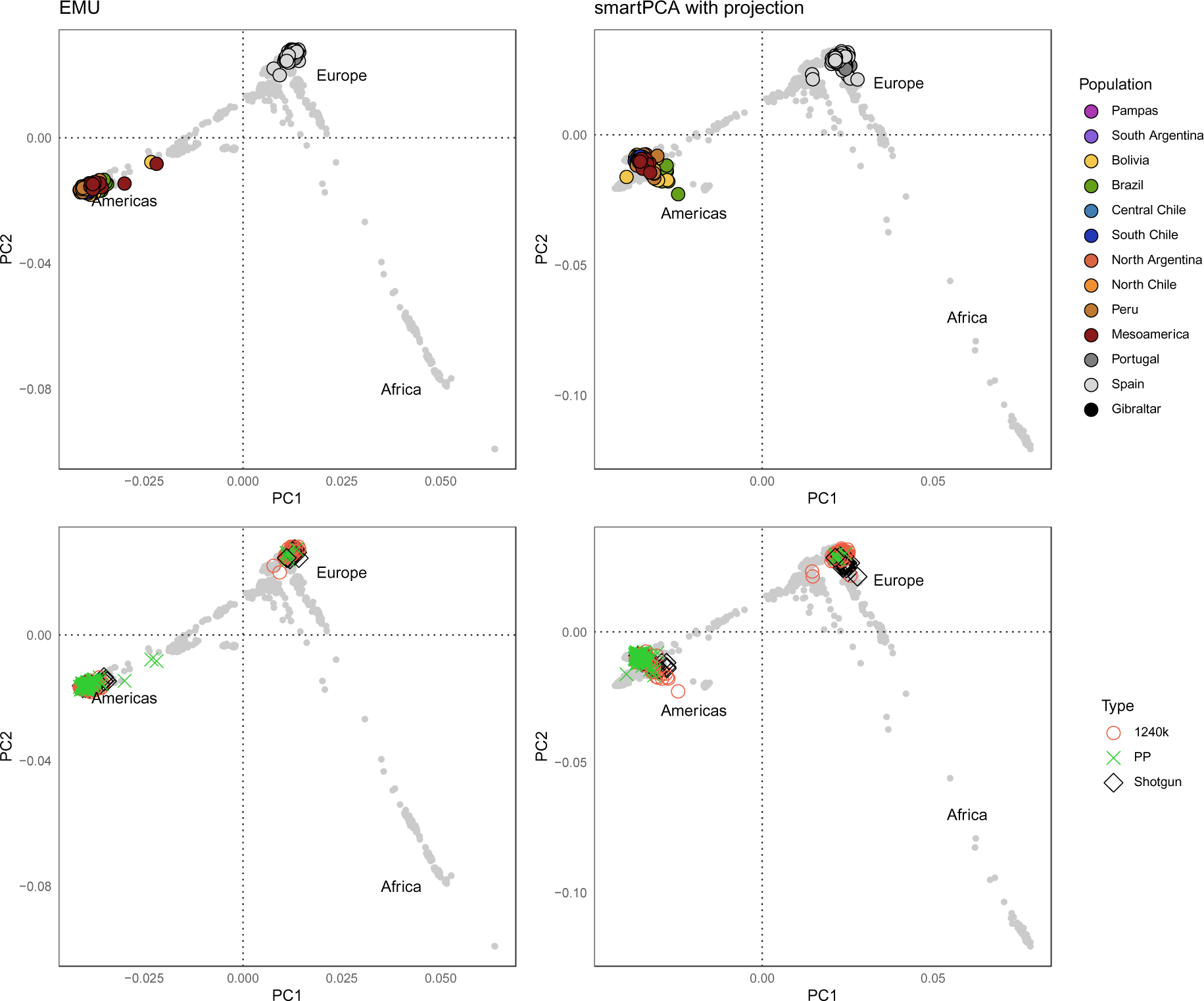
PCAs computed with EMU (left) and smartPCA (right) of ancient genomes from South America, Central America, and Iberia coloured by region (top row) or data type (bottom row). Ancient samples are projected by smartPCA onto the modern diversity of the HGDP shown in grey (Bergström et al., 2020).

### Genetic relatedness

We ran READ for pairwise combinations of four individuals from Peru for which we had generated shotgun data and single round PP-enriched data as a part of Experiment B (see Methods). These individuals represent the only samples for which both shotgun data and PP-enriched data were available. Given that only a single round of enrichment was used in these PP assays, technical bias is expected to be reduced relative to double-round enrichments.

The four artefactual “true twins” pairs (i.e., the same individual assayed with different methods; e.g., IndA.PP.1rnd.B and IndA.Shotgun) were the most related of all combinations with all average P0 values <0.4 (Fig S1). The remaining “non-twin” pairs all had a P0 value of close to 0.5, with small increments between them. Notably, PP-enriched pairs (e.g., Ind1.PP.1rnd.B and Ind2.PP.1rnd.B) were always more closely related (lower P0) than the respective shotgun pairing (e.g., Ind1.Shotgun and Ind2.Shotgun) (Fig S1). This suggests that the PP bias effects READ inferences, although it is subtle in this instance.

### Comparison of strategies to reduce bias

We conducted two wet-lab experiments with altered enrichment protocols and applied a total of 18 bioinformatic strategies, aiming to find a strategy that best reduced the Prime Plus bias. All strategies are detailed in Table S5. Bioinformatic strategies are categorised according to whether they 1) reduce mapping reference bias, 2) reduce the impact of ancient DNA damage, 3) remove specific “biased” SNPs, 4) employ statistical filters based on allele frequencies or 5) call genotypes with alternate methods. Our test population comprised the ancient Peruvian samples, the only population for which we generated both Prime Plus and shotgun data (Bongers et al., 2020). This allowed us to compare the effect of bias-reduction strategies in an ƒ_4_ test using statistics of the form ƒ_4_(Mbuti, Spain.PP.2rnd; Target, Peru.Shotgun), where the target rotates between each of the different bias reduction strategies applied to the Peruvian population. We chose Spain.PP.2rnd for this assessment for the following reasons: i) the Spanish dataset was generated in a different lab, by different individuals, and was sequenced on a different sequencing machine, than the Peruvian dataset (Table S2), thereby ruling out batch effects and isolating the Prime Plus enrichment kit as the underlying cause of any bias; and ii) the Spanish and Peruvian populations are sufficiently divergent and lack evidence of recent admixture (the Peruvian data date to pre-colonial times) such that the impact of Prime Plus bias is readily observable, but is not stronger than the population affinities, as would be the case if we were comparing two more closely related groups (such as populations from the same continent). To quantify the impact of Prime Plus bias relative to population affinity between the Spanish and Peruvian samples, we included Spain.1240k and Spain.Shotgun from publicly available data as target populations.

In total, we calculated ƒ_4_ statistics in the following four configurations to robustly estimate the Prime Plus bias and determine the best strategies to reduce it (Fig 4):

1) ƒ_4_(Mbuti, Spain.PP.2rnd; Target, Peru.Shotgun)
2) ƒ_4_(Mbuti, Spain.PP.2rnd; Target, Peru.1240k)
3) ƒ_4_(Mbuti, Spain.1240k; Target, Peru.Shotgun)
4) ƒ_4_(Mbuti, Spain.1240k; Target, Peru.1240k)

**Figure 4:**
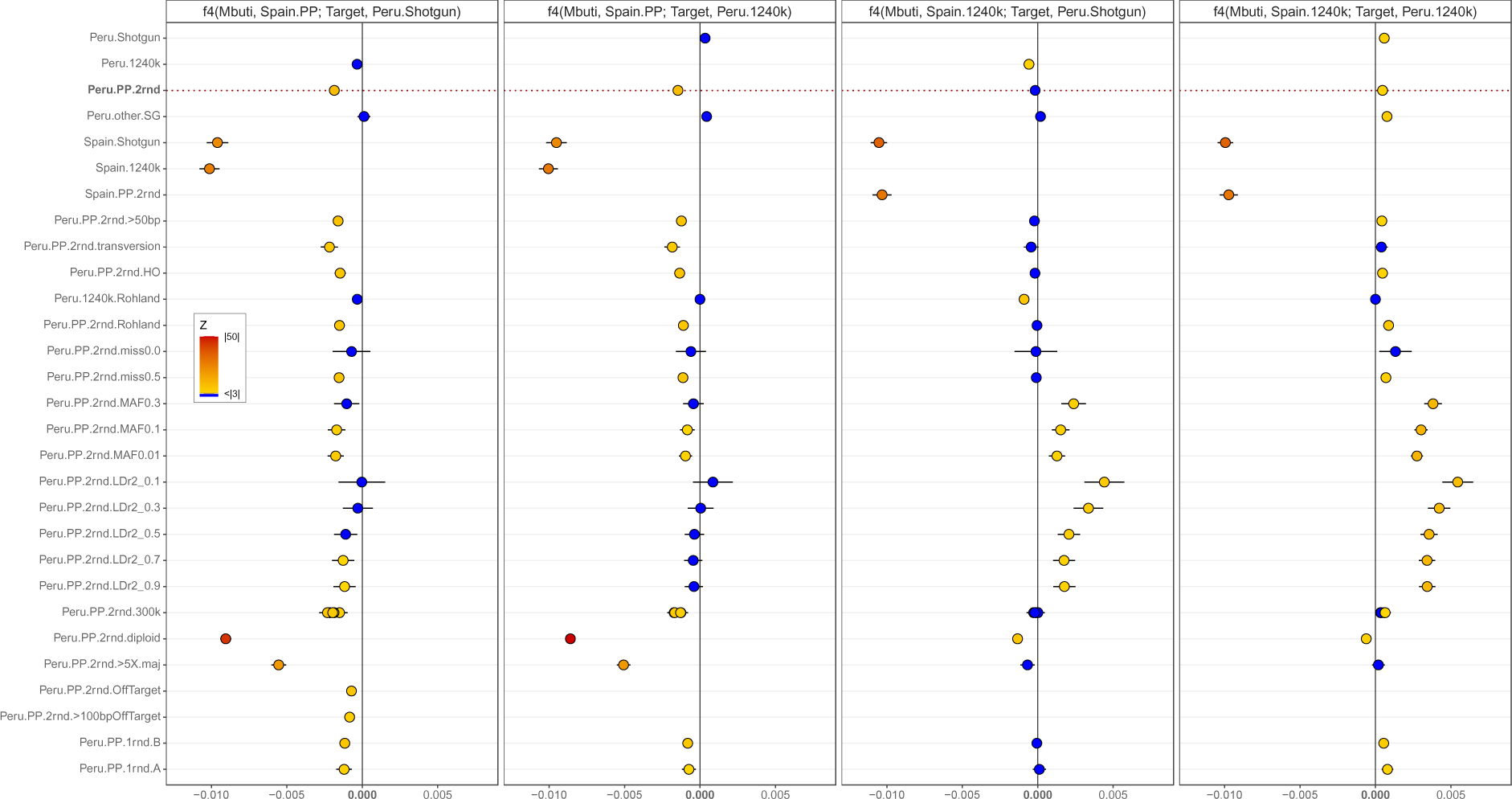
ƒ_4_ statistics showing the effectiveness of 20 bias-reduction strategies. Test configurations are, from left to right: ƒ_4_(Mbuti, Spain.PP.2rnd; Target, Peru.Shotgun), ƒ_4_(Mbuti, Spain.PP.2rnd; Target, Peru.1240k), ƒ_4_(Mbuti, Spain.1240k; Target, Peru.Shotgun), ƒ_4_(Mbuti, Spain.1240k; Target, Peru.1240k). “Target” groups are shown along the y-axis, being either one of the 29 strategies used to resolve the Prime Plus bias, or shotgun or 1240k data from Peru or Spain. Z-scores and the number of SNPs used in each test are recorded in Table S6. Non-significant *Z*-scores (|*Z*| < 3) are shown in blue, and significant *Z*-scores (|*Z*| > 3) are indicated with a yellow-red gradient. Error bars represent ± 2 se. The vertical dashed line represents the observed ƒ_4_ statistic in each configuration when Peru.PP.2rnd was the Target group. Note that bias-reduction strategies are only applied to the target population and the ƒ_4_ statistic is calculated from SNPs overlapping between the target and the other populations that remain unchanged between tests. Note that in the ƒ_4_ statistics where the target population is off-target or out-of-range genotypes (Peru.PP.2rnd.OffTarget, Peru.PP.2rnd.>100bpOffTarget), all four populations were genotyped in the respective alternative way to ensure overlapping SNPs, and therefore these are not exactly comparable.

The reason for using the first two configurations is that by comparing the target populations with Peruvian shotgun data, we are measuring our attempts to resolve the Prime Plus bias against the least biased data currently available. However, this may be an unreasonable standard to meet, given that it is known that legacy 1240k data also has some biases (Rohland et al., 2022), and the bulk of published ancient DNA data is legacy 1240k data. Therefore, while this comparison may not be a perfect estimation of total bias reduction, it will provide more useful insights for researchers given that most projects will leverage large proportions of 1240k reference data. Following the same logic, in the third and fourth configurations, we also computed ƒ_4_ statistics with Spanish legacy 1240k data (Fig 4), to gain a deeper understanding of the merits of alternate strategies for reducing bias in different combinations of PP, legacy 1240k and shotgun data. The full list of ƒ_4_ statistics computed in this study are detailed in Table S6.

As a positive control, we included a different set of ancient shotgun genomes from Peru (Peru.other.Shotgun) as a Target population, which resulted in ƒ_4_ results that were not significantly different from zero (Z < 3), as expected. The main aim of this process was to identify a strategy that would behave like the positive control, i.e., that would reduce the enrichment bias (Peru.PP.2rnd; dashed vertical line, Fig 4) so the ƒ_4_ values are not significantly different from 0. In the first configuration ƒ_4_(Mbuti, Spain.PP.2rnd; Target, Peru.Shotgun), a result not statistically different from 0 would indicate the bias was alleviated, and in the second configuration ƒ_4_(Mbuti, Spain.PP.2rnd; Target, Peru.1240k), the same result would suggest that the PP data is at least co-analysable with 1240k data under specific configurations. Importantly, the third configuration ƒ_4_(Mbuti, Spain.1240k; Peru.PP.2rnd, Peru.Shotgun) (Fig 4) resulted in an ƒ_4_ result that was not significantly different from zero, indicating that there was no significant bias between these PP and legacy 1240k datasets. Finally, for the fourth configuration ƒ_4_(Mbuti, Spain.1240k; Peru.PP.2rnd, Peru.1240k) (Fig 4) the ƒ_4_ result is significantly positive, providing further evidence for the previously reported bias in the legacy 1240k enrichment (Rohland et al., 2022). Therefore, the purpose of including Spain.1240k among our tested configurations is to find a bias-reduction strategy that does not significantly impact these ƒ_4_ statistics, and that allows the confident application of bias reduction strategies to different combinations of the Prime Plus and other data types and under alternate ƒ_4_ statistic configurations.

#### Ancient DNA damage or reference bias

The filter for minimum read length (Peru.PP.2rnd.>50bp) suggested by (Günther & Nettelblad, 2019) to reduce mapping reference bias had minimal effect on the ƒ_4_ statistics, suggesting that the observed assay bias is not strongly impacted by very short reads.

Filtering out transition SNPs (A/G and C/T), thus removing any possible ancient DNA damage (Peru.PP.2rnd.transversion) had a negligible effect on the ƒ_4_ statistics scores. Indeed, because all libraries were half-UDG treated and the ends of sequencing reads trimmed, we did not anticipate that ancient DNA damage would have a large impact.

#### SNP-set based filters

Strategies to reduce bias by removing certain SNPs included filtering to include only the Human Origins SNP set (Peru.PP.2rnd.HO; 593,120 SNPs retained) and a filter previously reported (Rohland et al., 2022) to enable the co-analysis of legacy 1240k and Prime Plus data (Peru.PP.2rnd.Rohland; 803,662 SNPs retained). Retaining only the Human Origins SNPs had minimal effect on the ƒ_4_ results. Notably, the application of the Rohland filter did not alleviate bias in any of the configurations of the ƒ_4_ statistics that we tested. In some cases application of the Rohland filter resulted in the ƒ_4_ values shifting slightly toward zero, but in other cases slightly away, which is likely due to an increase in uncertainty from the approximately 2-fold reduction in the number of SNPs. Overall, this indicates the Rohland filter does not work as reported on the present dataset, suggesting that it is not a universally applicable solution for capture bias.

#### Allele frequency-based filters

We further examined the bias reduction efficacy of a series of filters based on allele frequencies, including filters for SNP missingness, minor allele frequency and linkage disequilibrium. Of the SNP missingness filters, a strategy that excluded sites missing in >50% of individuals (Peru.PP.2rnd.miss0.5) did not notably shift the ƒ_4_ value towards zero. However, excluding all SNPs exhibiting any missingness (Peru.PP.2rnd.miss0) did move the value toward zero, but it also reduced the number of SNPs so drastically that this strategy would probably not be suitable for most paleogenomic studies. For SNP filters based on minor allele frequency (Peru.PP.2rnd.MAF0.3, Peru.PP.2rnd.MAF0.1 and Peru.PP.2rnd.MAF0.01), ƒ_4_ results were shifted toward zero as the stringency of the filter increased, but this also led to increasingly large reductions in the number of analysable SNPs. Interestingly, in the ƒ_4_ configurations with Spain.1240k, the minor allele frequency filters produced positive ƒ_4_ values indicating a bias towards the Peruvian data regardless of whether shotgun or legacy 1240k are used.

When applying LD-based filters, we identified that the shared drift with Spain.PP.2rnd reduces as the stringency of the filter increases, though this once again came at the cost of increasingly fewer retained SNPs. In the configuration ƒ_4_(Mbuti, Spain.PP.2rnd; Target, Peru.Shotgun), the ƒ_4_ result for *r*^2^ = 0.1 was very close to 0 (Peru.PP.2rnd.LDr0.1). Filters for less stringent *r*^2^ values filtered out fewer SNPs but did not improve ƒ_4_ results to the same degree (Fig 4). In the configuration ƒ_4_(Mbuti, Spain.PP; Target, Peru.1240k), the less stringent *r*^2^ = 0.3 produces an ƒ_4_ value not significantly different from 0, with the retention of more SNPs than the *r*^2^ = 0.1 filter. The use of this filter to alleviate PP bias could be considered by users in specific cases with high sequencing depth to accommodate the expected SNP loss. Randomly downsampling the Peru.PP data to an equivalent number of SNPs (300,000) yielded ƒ_4_ values with additional variance around the value of the unfiltered Peru.PP data (dashed lines, Fig 4) and higher absolute *Z*-scores (Fig 4). This suggests that the bias may in fact be creating artificial “linkage” among neighbouring SNPs, but that sufficiently stringent LD pruning to “correct” the value to zero requires discarding the majority of targeted SNPs. Accordingly, the value of linkage disequilibrium pruning may be considered as a potential solution to the PP bias on a case-by-case basis, with the caveat that improved bias correction will be offset to some degree by the loss of precision from reduced statistical power, and consideration will need to be given to the effect of LD pruning on the results of various analyses.

#### Alternative genotyping

We considered several genotyping approaches alternative to the standard random pseudo-haploidisation, including calling random diploid genotypes, using majority call pseudo-haploid genotypes at sites with > 5X coverage, genotyping off-target SNPs, and genotyping ‘out-of-range’ SNPs that were > 100 bp from the target sites. Calling off-target genotypes (Peru.PP.2rnd.OffTarget) and out-of-range genotypes (Peru.PP.2rnd.>100bpOffTarget) both resulted in a small shift of the ƒ_4_ values toward zero. Majority call genotypes (Peru.PP.2rnd.>5X.maj) behaved very differently between configurations. In configurations with Spain.1240k, the filter had a minimal effect, but with Spain.PP.2rnd, the majority call genotyping drastically shifted the ƒ_4_ results towards negative values, indicating that this approach increases the artefactual “shared drift” between PP data from Spain and Peru. Random diploid genotyping produced an ƒ_4_ value that was even more biased (Peru.PP.2rnd.diploid), mildly so when compared with Spain.1240k, but much more strongly for Spain.PP.2rnd. In fact, the latter value is close to the ƒ_4_ result where the target dataset is Spain.Shotgun. This means that using random diploid genotype calls exacerbates the artefactual shared drift driven by the PP enrichment bias to approximately the same magnitude as true shared drift between Spanish populations (Fig 4).

#### Laboratory experiments

In our final set of bias reduction strategies, two non-standard lab experiment protocols that used a single round of enrichment (Peru.PP.1rnd.A and Peru.PP.1rnd.B) both led to minor bias reductions. Protocol B, which lowered the hybridisation temperature performed slightly better than Protocol A. While the overall improvement exhibited by both methods was minor, the bias remained considerably greater than the bias observed in the legacy 1240k data (Peru.1240k).

### Allele calls

We observe strong GC bias amongst the data generated with the legacy 1240k baits, whereas the PP data display comparatively lower levels of bias that are broadly comparable with shotgun datasets (Fig 5). It is important to note that the bias apparent in the legacy 1240k data was not a concern prior to the release of alternate enrichment kits, simply because all the data generated using the legacy 1240k baits are comparable. We observe that across all three methods, the GC bias is stronger for transitions—i.e., allele pairs A/G and C/T where both alleles are either purines or pyrimidines, respectively— which represent the majority of target sites (80%). Though our understanding of the biochemical mechanisms underlying the PP bias is limited by a lack of knowledge of the bait design, our data clearly demonstrate a systematic difference in the pattern of allelic biases between the legacy 1240k and PP assays.

**Figure 5:**
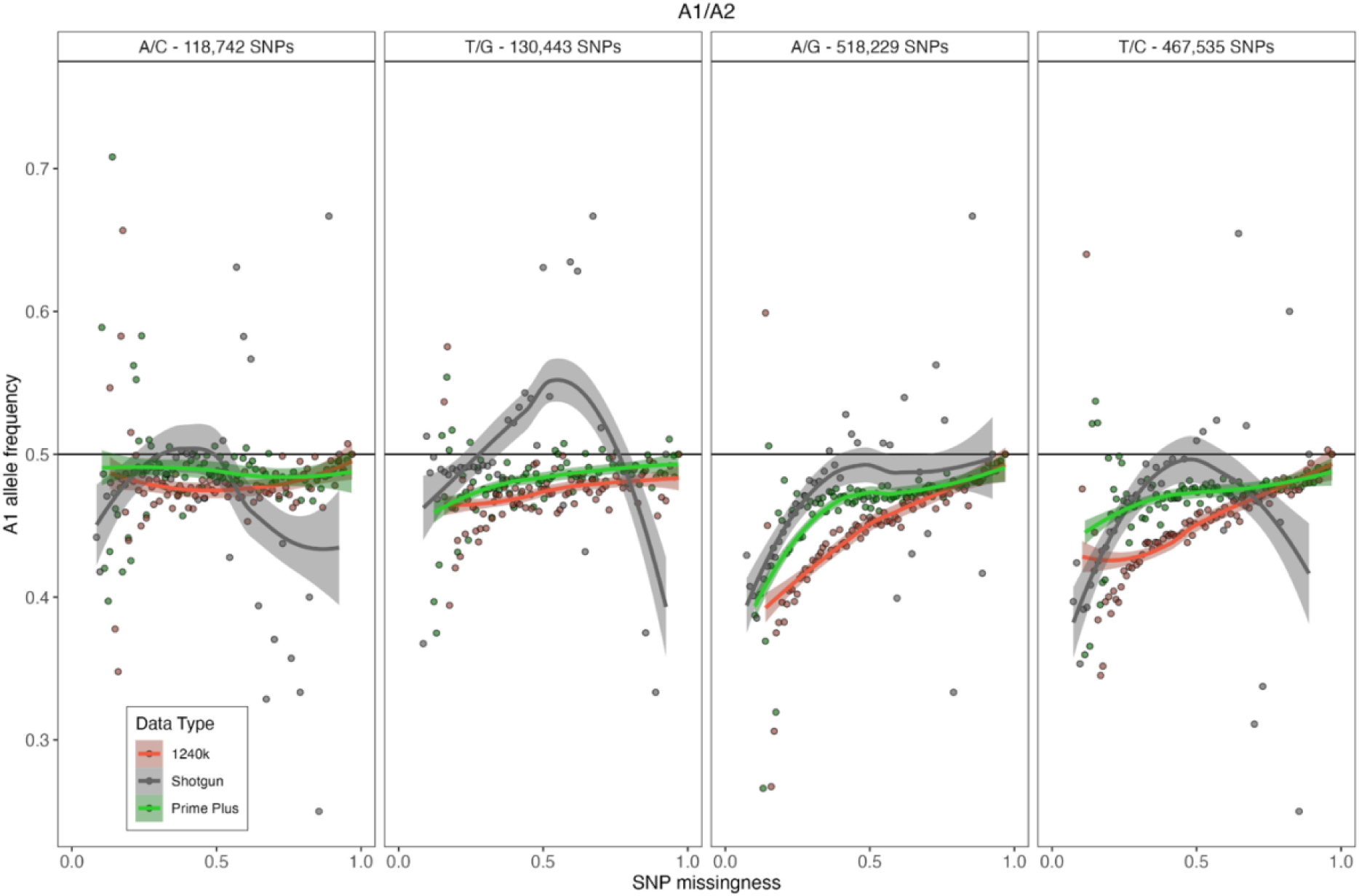
Mean frequency of A1 allele calls in different data types as a function of SNP missingness. Results for the four different A1 allele pairs are provided in separate panels, with inset data points showing the mean allele frequencies calculated across all individuals in successive SNP missingness bins. SNPs that are fixed within each dataset were removed to avoid skewing the mean A1 calls. The header of each panel shows the number of target SNPs of the corresponding A1/A2 pair observed in the Prime Plus assay.

## DISCUSSION

Results presented in this study support the previous report of a bias when enriching SNPs at ∼1240k sites using the Daicel Arbor Biosciences Expert Human Affinities Prime Plus enrichment kit, and extend the previous findings to ancient Indigenous American (i.e., non-Eurasian) populations. In accordance with (Rohland et al., 2022) we find that the biased data strongly affect ƒ-statistic calculations, where the shared genetic drift (i.e., the evolutionary history) between groups is explicitly estimated. Note that we consider the effect strong, not because of the magnitude of any quantitative measurements, but due to the corruption of expected demographic relationships between populations that is driven by this bias. In accordance with results reported by (Rohland et al., 2022), we find that the assay bias does not strongly affect PCA at least in the first three prinicipal components. Additionally, for the first time, we observe a (in this case mild) bias effect on the predicted relatedness between individuals using the ancient DNA relatedness test READ.

### Batch effects

The initial observation that PP data had an unexpected excess of shared drift with other PP-generated data in ƒ-statistics prompted us to investigate if this artefact was due to a batch effect arising from certain samples, specific labs having done the work, DNA sequencing machines, or resulted from the distinct demographic history of populations in question. We can confidently rule out each of these hypotheses, as the lab work was performed on numerous sample sets, in different labs, by multiple individuals, sequenced in different facilities using different sequencers (Table S2), with the only common factor to all biased samples being the PP enrichment. Our outgroup-ƒ_3_ statistics calculated between European and South American populations (Fig 1) confirmed that the bias affected these two highly diverged populations suggesting that it is most likely a systematic bias that affects all human ancestries. However, we did not have access to the appropriate shotgun data to determine if the enrichment bias has differential effects by ancestry source, and this hypothesis requires additional testing.

### Describing the Prime Plus bias

#### PCA

We performed principal components analysis in several ways with the aim of addressing multiple analytical approaches that researchers might take. To accommodate differing geographical scales, we computed separate PCAs at continental and global scales. While the approach of using smartPCA with projection has long been considered the gold standard, it should be noted that this method works best with high-quality ancient samples that have low missingness and a well-developed reference database of modern diversity that captures the genetic variation present in the ancient populations, therefore enabling the most accurate projection. Hence, the PCA published by Rohland et al. 2022 could be expected to show good results as they projected 15 high-quality ancient genomes onto the modern diversity of western Eurasia, the most studied region of the world. However, this is generally less achievable in understudied regions where modern reference data is lacking and/or when working with low-quality samples. This is what motivated the simultaneous use of EMU (designed to handle data missingness and does not use projection) (Meisner et al., 2021) and smartPCA both with and without projection onto modern diversity.

At both the the continental and global scale, both EMU and smartPCA without projection produced PCAs that reflects South American population structure in accordance with previous results <u>(Posth et al. 2018; Nakatsuka et al. 2020; Nakatsuka et al. 2020; Martiniano et al. 2017; Olalde et al. 2019; Valdiosera et al. 2018)</u> and patterns suggesting assay bias are not evident within the first three principal components (Fig 2). However, we caution that the results of smartPCA are variable by the version used and the data inputted, therefore these results do not preclude the emergence of assay bias in a lower principal component or within a different dataset.

At a global scale, both smartPCA with projection onto the modern diversity of the HGDP data (Bergström et al., 2020) and EMU performed similarly. Note that EMU does not have projection functionality and instead includes all individuals in the calculation of eigenvectors while imputing sites with missing data (Meisner et al., 2021). There does not seem to be any obvious assay bias in either PCA output, although this might still be present in a lower-level PC. Furthermore, discrepancies between the PCA methods, and the effect of assay bias may become more impactful on results and interpretations at finer-scales.

Overall, we suggest constructing PCAs using smartPCA, projecting ancient genomes onto modern diversity where possible. However, in the case that modern reference data is lacking, PCA plots from smartPCA and EMU should be produced and compared, especially in cases where PP data, or data with high rates of missingness are included.

#### Genetic relatedness

We assessed the effect of PP bias on results from the genetic relatedness estimation software READ (Monroy Kuhn et al., 2018). We found that pairwise estimations between PP libraries were always more closely related than shotgun library pairs of the same four samples (Fig S1). Given that these PP libraries were only enriched for a single round, we would expect this bias to be even stronger in PP libraries that follow the standard two-round enrichment protocol (reflecting patterns observed in the ƒ_4_ statistics; Fig 4). In the present case, the bias is subtle and unlikely to lead to, for example, a false-positive first-degree relationship from an unrelated pair. While our ability to make more general conclusions is clearly limited by the sample set, our results tentatively suggest that READ inferences should be reasonably robust if run on a dataset of only PP data, but users should exhibit caution when combining different data types.

#### ƒ-statistics

Our analyses of the PP bias in both outgroup-ƒ_3_ and ƒ_4_ statistics showed that the bias is strong enough to corrupt expected relationships between populations from the same continent (Fig 1). We reproduced the legacy 1240k bias that has been previously reported (Rohland et al., 2022) and showed that it is relatively subtle compared to the PP bias, indicating that it is less impactful on population genetic inferences.

Importantly, our analyses suggest that it should be possible to use Prime Plus data in a comparative outgroup-ƒ_3_ if the target population is the only one enriched with Prime Plus, that is statistics of the form ƒ_3_(Outgroup.notPP; target.PP, X.notPP) where both the outgroup and population X are not enriched with Prime Plus. This is suggested by our ƒ_3_ statistics (Fig 1A) where Peru.PP.2rnd is the Target population, as the other non.PP South American populations reproduce the expected relative relationships previously reported (Nakatsuka, Lazaridis, et al., 2020; Nakatsuka, Luisi, et al., 2020; Posth et al., 2018) despite none having been enriched with the Prime Plus baits. However, the limitation is that a population enriched with Prime Plus could not be split into subpopulations, such as Peru_A.PP.2rnd and Peru_B.PP.2rnd, and rotated in the X population position, or in scenarios where one is X and the other Test, but then compared to ƒ_3_ statistics of a different form. Presumably, an ƒ-statistic would also be viable if all X groups were PP-enriched because the enrichment bias would be shared by all data and should not corrupt demographic interpretations of the results. See Table S4 for a comprehensive list of acceptable outgroup-ƒ_3_ and ƒ_4_ statistic forms and, though not directly investigated here, using the same logic we also list qpWave and qpAdm configurations and their relative reliability under PP bias.

### Strategies to reduce bias

The bias observed in ƒ-statistics is both persistent and systematic, which allowed us to test a total of 20 experimental and analytical protocols to characterise the bias and alleviate its disruptive impact on inferences from common population genetic analyses. Many of these analytical protocols produced marginal improvements, but no protocol was sufficient to remove all biases (*Z*-score remained such that |*Z*| > 3, and/or resulted in low SNP retention).

#### Ancient DNA damage or reference bias

Neither ancient DNA damage nor reference bias appeared to contribute to the bias, as removing transition sites (all SNPs that could potentially be ancient DNA damage) and removing reads shorter than 50 bp had little impact on bias in the ƒ_4_ statistics (Fig 4).

#### SNP-set based filters

Next, we investigated if a subset of the 1240k SNPs was driving the bias and could therefore be removed to allow safe co-analysis of PP enriched samples with other data types. This idea was initially suggested by Rohland and colleagues (Rohland et al., 2022), who published a list of SNPs that they suggested should be suitable for co-analyses of shotgun, legacy 1240k, Twist, and PP data. However, our tests of the Rohland et al. filter produced no notable improvement on the ƒ_4_ statistics when co-analysed with 1240k or PP data (Peru.PP.2rnd.Rohland, Fig 4). Note that to build this filter Rohland and colleagues looked for improved *Z*-scores, i.e., closer to zero, but did not report the ƒ_4_ statistics that are informative to estimate the effect on population relatedness. When using any SNP filter, the removal of SNPs will likely increase the standard error of the ƒ_4_ statistic, leading to decreased statistical power and precision. However, it is not possible to predict the deviation from the true ƒ_4_ statistic. Therefore *Z*-scores alone do not serve as a measure of the bias-reduction of a filtering strategy. This emphasises the importance of testing hypotheses in the context of the study and understanding the expected relationships of the populations in question.

Subsetting our data to include only the ∼600k SNPs in the Human Origins set had a similar effect to the Rohland filter. This may be related to the overlap between the two sets, 61% of the Human Origins SNPs are also present in the Rohland filter set. (367,332 intersecting SNPs).

In summary, none of these filtering strategies corrected the PP bias satisfactorily. Furthermore, while Rohland et al. acknowledge there is room for improvement in their filtering approach, it is possible that their filter is well-calibrated for the Eurasian ancestries present in their dataset but does not represent a universal filter to mitigate Prime Plus capture bias in all ancestries. In any case, it is less than ideal to have enrichment kits on the market that would require discarding a subset of target SNPs when combining multiple data types. This will negatively affect not only users of the Daicel Arbor Biosciences Human Affinities kits — and potentially other commercial kits — but also impact subsequent analyses by other researchers that incorporate published PP datasets.

#### Alternative genotyping

Genotype calling in paleogenomic studies typically involves randomly sampling one read covering a targeted SNP site, known as ‘pseudo-haploidisation’, making it possible to generate allele calls at sites with low coverage (Morris et al., 2011). Given that our experiments with SNP filtering did not produce a universal solution and following the assumption that every targeted SNP site may be affected by the bias, we investigated the efficacy of alternate genotype calling for bias reduction. In our first assessment, we called pseudo-haploid genotypes at off-target SNPs, which we hypothesised would be equivalent to shotgun data and therefore unbiased if the bias was driven by the bait design. Similarly, we also called random pseudo-haploid genotypes at sites more than 100 bp away from a target locus to mitigate the possibility of an “off-target” site being covered by a probe that had correctly hybridised to a nearby target locus. The ƒ_4_ statistic for both off-target and out-of-range genotypes did improve compared to the on-target results, producing values that were not significantly different from zero (Fig 4). Interestingly, the ƒ_4_ statistic for the out-of-range genotypes was not notably different from the off-target estimation. Given that all non-bound DNA fragments are theoretically washed away during the enrichment protocol, the off-target bait bias is likely due to an as-yet-unknown biochemical bias that systematically affects bait binding to DNA at non-target locations, but it is unlikely this is caused by on-target hybridised baits covering nearby SNPs within a 100 bp range.

Additionally, we used an alternate pseudo-haploid calling approach which requires target SNPs to have a higher minimum coverage (5X) and takes the allele represented by the majority of reads as the final call. This experiment (Peru.PP.2rnd.>5X.maj, Fig 4) yielded a dramatic shift of the ƒ_4_ results towards large negative values, further exacerbating the artefactual shared drift and bias. This is strong evidence that the bias is conferred by the bait design, and results in the persistent overrepresentation of one allele over the other at heterozygous sites. Concordantly, our random diploid genotype test produced even more biased ƒ_4_ statistics, such that the estimated levels of shared drift observed between the Spanish PP data and Peruvian PP data that used random diploid genotype calls is of a similar magnitude to the true shared drift between Spanish PP generated and Spanish shotgun data. This is evidence of a remarkably strong systematic bias that has the potential to corrupt all paleogenomic studies combining PP with other data types if not handled carefully. These two results are strongly suggestive of a systematic allelic bias that impacts all heterozygous SNPs and is potentially caused by the molecular design of the PP baits.

#### Laboratory protocol experiments

Our two experiments with single-round PP enrichment yielded ƒ_4_ values closer to zero than those produced by the standard two-round enrichment. This improvement is likely because whatever bias occurs during the first round of bait–DNA binding becomes amplified during the second hybridisation step because it is acting on an already biased set of DNA fragments. This interpretation is supported by the sequencing statistics (Fig S3) showing that repeated capture rounds result in an increased endogenous DNA percentage but decreased library complexity (observed as fewer unique reads when sequenced). Additionally, any biased sampling that has occurred during the initial binding will be exponentially amplified during the PCR step, so we would expect a reduction in PCR cycles to improve this. The reduced hybridisation temperature should also decrease the specificity of hybridisation and thus reduce allelic bias, but simultaneously increase off-target binding, thus reducing the percentage of endogenous DNA enriched.

### Characterising the bias

Given the evidence that the bias affects all targeted SNPs, we investigated whether this effect was independent from the SNP transition, hoping that we might be able to apply some relatively simple statistical scaling of the genotype calls that would provide co-analysable genotypes. However, our investigations of allelic bias (Fig 5) clearly show that the capture bias produces complex effects on allele calls. We can only confidently observe that the pattern of GC bias in PP data is different to the 1240k and shotgun data. Further, the patterns of missingness evident in the PCA analyses consistently differ between data types (Fig S2), indicating that there may be several layers of biochemical bias at play including that each assay preferentially captures certain SNPs, and/or favours the capture of different alleles at heterozygous sites, with allelic preference being more exacerbated in PP data.

### Study limitations

The main limitation of this study is that we did not have samples for which data was generated in all of the three methods addressed. This is due to the opportunistic nature of the study in that we collated this dataset when the PP bias was initially observed, after the data generation process. However, we have made every effort to combat this limitation by maximising the size of the dataset and explicitly stating assumptions where we’ve made them.

In our READ analyses we were limited to a dataset of only four individuals for which we had shotgun and PP data, and a further limitation is that this PP data was from one of our single-round enrichment experiments and thus does not fully represent the PP bias under investigation. To further understand the impact on READ, both shotgun and two-round PP enriched data would need to be generated for a larger number of samples, from a more diverse set of relationships to test the impact of bias on truly related individuals.

The main outstanding hypothesis to investigate is whether this bias has differential effects on different ancestries. This would require a large dataset of PP and shotgun data from the same individuals across more ancestries than in the present study. While an interesting question, we believe this study presents sufficient evidence that the PP assay is biased.

### Implications

For the past decade in-solution hybridisation enrichment has been a go-to method for obtaining affordable, high-quality, and informative data for paleogenomic studies. However, irrespective of the bait design, the biochemistry of DNA hybridisation means that one bait will always have a higher binding affinity with its targeted allele than any other. Greater differences in these affinities result in increased allelic biases in the generated datasets. This is evidenced by our assessment of alternative enrichment protocols, where the single-round enrichments have less endogenous DNA (Fig S3), but also appear to be less biased (Fig 4). Therefore, we hypothesise that the enrichment efficiency of the PP reagent is in direct trade-off with the biochemical allelic bias.

It is worth reiterating here that all capture methods used in paleogenomic studies will lead to ascertainment bias, meaning that one must carefully consider both the selection of methods and the use of comparative published data. Previously, the monopoly on data generation by labs with access to the legacy 1240k enrichment protocol somewhat insulated the community from this compatibility issue. However, now that data can be generated using different enrichment kits, researchers face a new set of challenges whenever they integrate data generated with new capture assays with previously published data generated using the legacy 1240k—the most comprehensive dataset to date that underpins many of the findings in paleogenomics. Additionally, researchers utilising the Twist reagent should benchmark the data similarly to the present study to establish its co-analysability with other data types.

Here, we have tested some of the most used analyses in paleogenomic studies, though not an exhaustive list. Principal Components Analysis is a standard way to understand population structure, and outgroup-ƒ_3_ and ƒ_4_ statistics, along with their more complex derivatives, are integral to reconstructing population relationships and evolutionary history, while READ is a standard for genetic relatedness analysis. We have found that PP-enriched data bias the results of READ and ƒ statistics when used in combination with other non-PP datasets, and therefore we caution potential PP users that analyses of such multi-assayed datasets may lead to erroneous conclusions if not interpreted carefully.

Having comparative shotgun data for a subset of samples in a study, or from samples expected to have a very similar ancestry should allow researchers to estimate and control for the PP bias in the specific methods they wish to utilise. Further, while we explicitly test PP data, we advise researchers to exhibit similar caution when using the Daicel Arbor Biosciences Expert Human Affinities “Complete” and “Ancestral Plus” enrichment kits, as the bait chemistry and enrichment protocol is the same as the PP kit, and only the number of target loci differ.

### Guidelines

- When calculating ƒ-statistics only compare Prime-Plus-generated data with other Prime-Plus-generated data; OR
- Only compare one Prime-Plus-generated target population/individual with other non-Prime-Plus-generated populations/individuals (See Table S4 for a full description).
- Beware that Prime-Plus-generated data attract each other and that this may artificially confirm/reject your hypothesis depending on the study design.
- Explicitly assess the impact of Prime-Plus bias in your data when running analyses that have not yet been investigated.
- Recommend to shotgun sequence a subset of enriched libraries as a control set if possible, ideally select libraries to represent a range of ancestries.
- Always annotate your data as Prime-Plus-generated when publishing, sharing with other researchers, or incorporating into any publicly accessible database.
- Always annotate the data type of both newly generated and published data used in your study so this can be factored into interpretations.
- For best results compute PCAs using smartPCA with projection where possible and otherwise with EMU to compare results; if clustering appears driven by capture assay, pursue alternative analyses.
- In general, pursue multiple lines of evidence to support your findings.

## Acknowledgements

We thank Adam Rohrlach for insightful manuscript revisions and assistance with statistical interpretation. We thank members of the Australian Centre for Ancient DNA’s Thesis Writing Group for their contributions to manuscript revisions, especially Olivia Johnson for assistance with R coding. We thank the members of the modern and ancient DNA research group at the Unitat d’Antropologia Biològica, Universitat Autònoma de Barcelona (Spain) for their contribution to data generation. We thank Linda R. Manzanilla and Grégory Pereira from the Instituto de Investigaciones Antropológicas, Universidad Nacional Autónoma de México (Mexico), and UMR 8096 Archéologie des Amériques, Centre National de la Recherche Scientifique (France), respectively, for their contribution to sample provision. LFS was supported by funding from the National Science Foundation (NSF - 1515138), RT by the Australian Research Council (DE190101069), BL by an Australian Research Council Future Fellowship (FT170100448).

## Author Contributions

R.D., B.L., L.F-S., M.P.W. and X.R-R. conceptualised the study. R.D., L.F-S., K.K., M.P.W. and X.R-R. performed laboratory work. R.D. and L.F-S. performed computational analyses. R.T., B.L., and L.F-S. funded the study. All authors contributed to manuscript writing and revisions.

## Data Accessibility and Benefit-Sharing Section

### Genomic Data

The newly generated genotype data in this study are available with DOI: 10.25909/24004665. We recommend against the inclusion of this data in other population genomic analyses due to the bias present.

### Code Availability

Code used for analyses and plot generation is available at https://github.com/roberta-davidson/Davidson_etal_2023-Capture-bias-paper.

