## Supplementary material for "Allelic bias when performing in-solution enrichment of ancient human DNA": Fig S

<u>Supplementary Information</u>

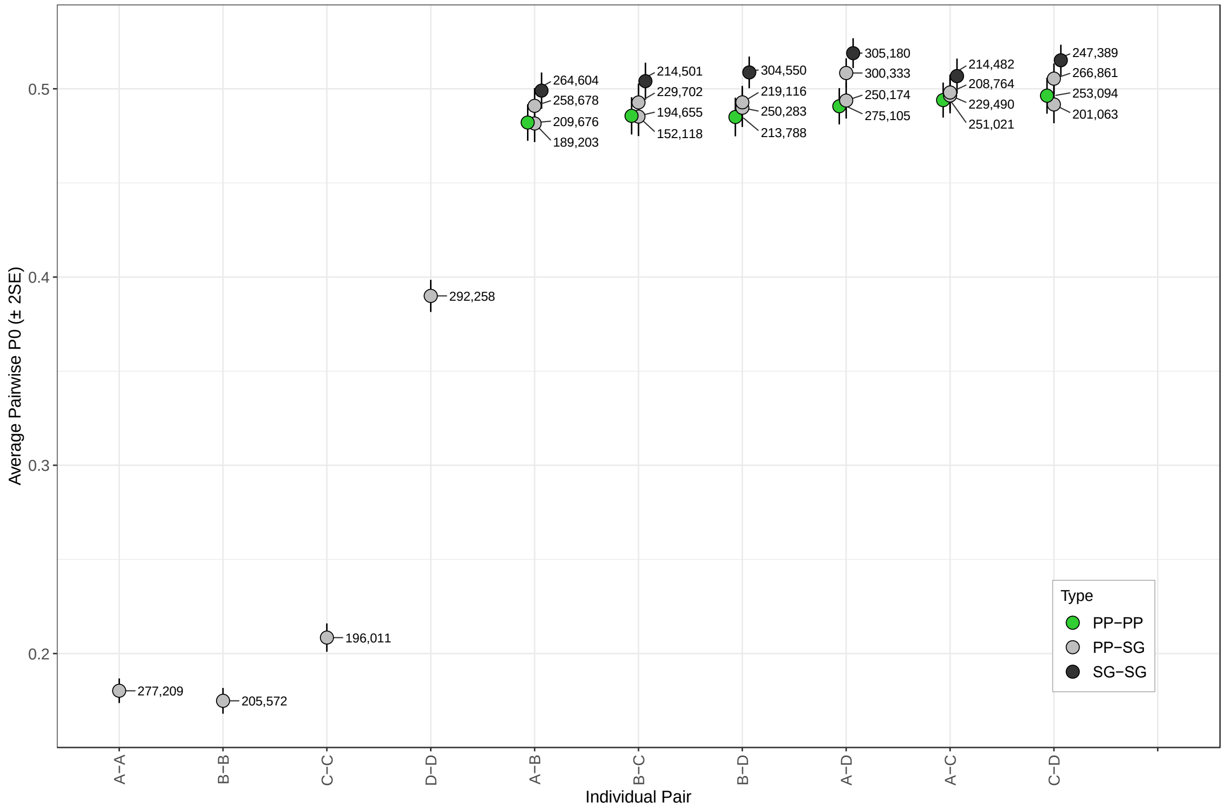

**Figure S1**: READ results for pairwise combinations of four individuals from Peru for which we had generated shotgun and single round Prime Plus enriched data as a part of Experiment B. Each point is labelled with the number of SNPs used in the pairwise comparison.

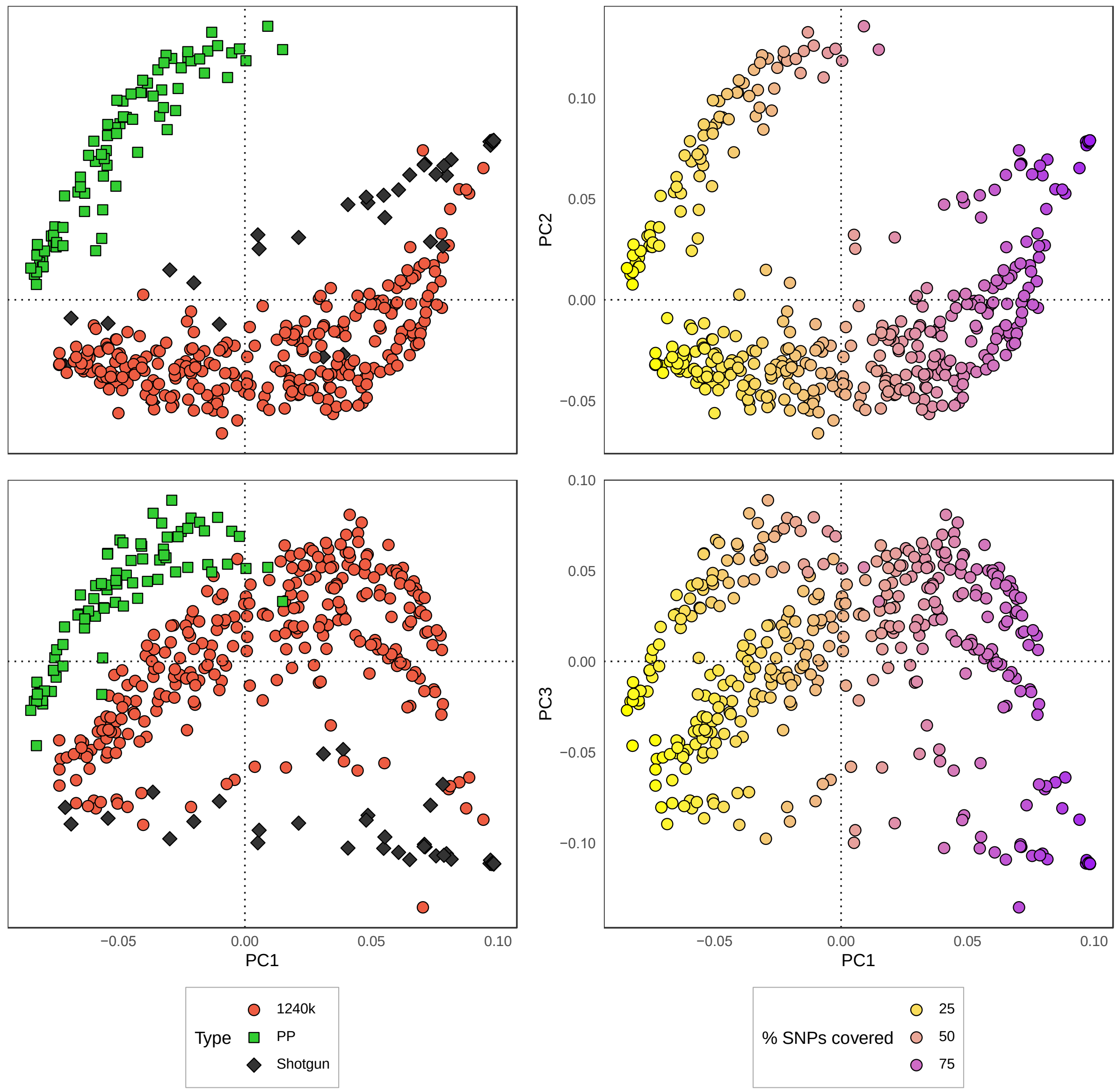

**Fig S2:** PCA of SNP missingness between data types (left column) computed with smartPCA. The right column is coloured by missingness frequency where yellow is high missingness and purple is low missingness.

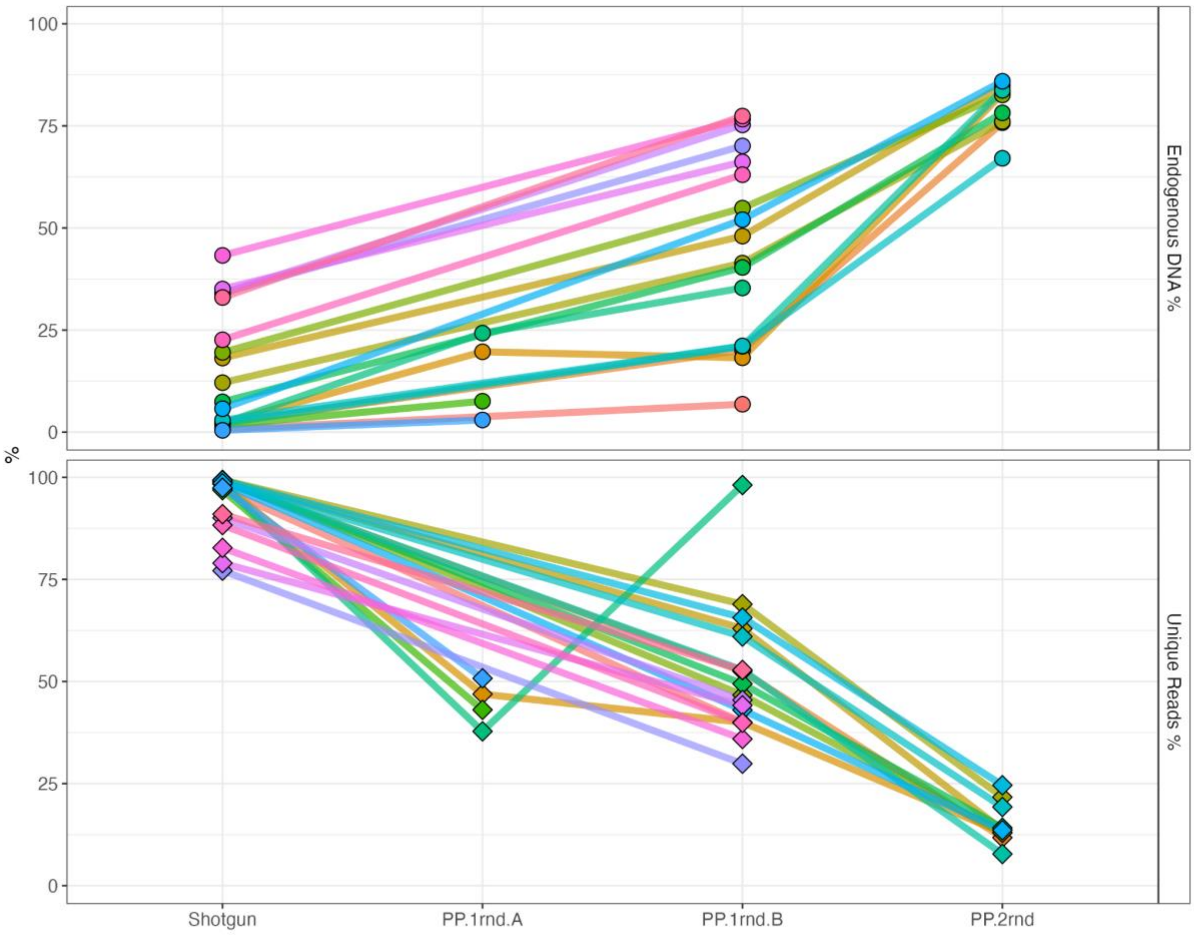

**Figure S3:** Key summary statistics from sequencing libraries at different levels of enrichment (x-axis), contrasting increasing endogenous DNA retrieved (top) with decreasing complexity, taken as % unique reads (bottom). Values are standardised by the total reads.

Tables S1-S3 and S6 are in the Supplementary Excel spreadsheet.

**Table S1:** Number of ancient samples per region for each data type represented in this study.

**Table S2:** Processing details of all Prime Plus enriched samples in this study.

**Table S3:** Metadata for all published samples used as comparative data in this study

**Table S4**: List of possible ways to structure outgroup-$f_3$ and $f_4$ statistics as well as qpAdm and qpWave configurations using PP data and their respective reliability.

| **$f_3$ statistic structure** | **Reliability and Expectation** |
|---|---|
| 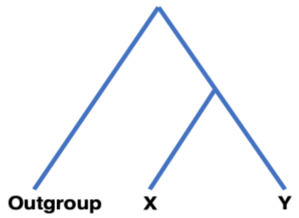  | *Where Outgroup is kept constant between comparisons and X and/or Y rotate between populations or individuals* |
| $f_3$ (Outgroup.PP; X.PP, Y.PP) | Reliable, bias affects all equally |
| $f_3$ (Outgroup.notPP; X.PP, Y.PP) | Reliable, under the assumption that bias does not significantly differentially affect different ancestries. |
| $f_3$ (Outgroup.notPP; X.sometimesPP, Y.sometimesPP) | Unreliable, X.PP will always have more shared drift with Y.PP than X.notPP. |
| $f_3$ (Outgroup.notPP; X.notPP, Y.PP) | Reliable, the bias in Target.PP is not shared with X or Outgroup so the test should be agnostic to it. |
| $f_3$ (Outgroup.notPP; X.notPP, Y.notPP) | Reliable when using shotgun or legacy 1240k, standard practice. |
| **$f_4$ statistic structure** | **Reliability and Expectation** |
| 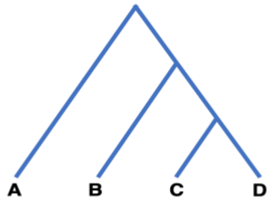  | *Where A is usually an outgroup population and B, C and D may rotate between populations or individuals* |
| $f_4$ (A.PP, B.PP; C.PP, D.PP) | Reliable, bias affects all equally |
| $f_4$ (A.notPP, B.PP; C.PP, D.PP) | Reliable, under the assumption that bias does not significantly differentially affect different ancestries. |
| $f_4$ (A.notPP, B.PP; C.notPP, D.PP) | Unreliable, B.PP and D.PP will be artificially attracted to each other |
| $f_4$ (A.notPP, B.notPP; C.PP, D.PP) | Reliable, as C and D share the PP bias, the test will reveal whether either received gene flow from B. |
| $f_4$ (A.notPP, B.PP; C.PP, D.notPP) | Unreliable, C.PP will be more attracted to the other PP population than it truly is. |

| | |
|---|---|
| $f_4$(A.notPP, B.notPP; C.sometimesPP, D.PP)<br>$f_4$(A.notPP, B.notPP; C.PP, D.sometimesPP) | Unreliable |
| $f_4$(A.PP, B.notPP; C.notPP, D.notPP)<br>$f_4$(A.notPP, B.PP; C.notPP, D.notPP)<br>$f_4$(A.notPP, B.notPP; C.PP, D.notPP)<br>$f_4$(A.notPP, B.notPP; C.notPP, D.PP) | Reliable, the bias in the one PP population is not shared with the others so the test should be agnostic to it. |
| $f_4$(A.notPP, B.notPP; C.notPP, D.notPP) | Reliable when using shotgun or legacy 1240k, standard practice |
| **qpAdm configuration** | **Reliability and Expectation** |
| *Where T = target, S = source, R = reference* | |
| (T) = T.PP, (S) = S.notPP, (R) = R.notPP | Reliable, under the assumption that the PP bias does not significantly differentially affect different ancestries or either of the shotgun and 1240k technologies. |
| (T) = T.PP, (S) = S.sometimesPP, (R) = R.notPP | Unreliable, bias in the matrix of $f_4$ statistics underlying qpAdm could lead to artificial relationships and weight estimations from the source to the target population. |
| (T) = T.PP, (S) = S.sometimesPP, (R) = R.sometimesPP | Unreliable, bias in the matrix of $f_4$ statistics could lead to incorrect estimation of the matrix rank in addition to the problems outlined above. |
| (T) = T.PP, (S) = S.notPP, (R) = R.sometimesPP | Unreliable, bias in the rank of the matrix of $f_4$ statistics can be driven by the artificial affinity between the T.PP target and R.PP right-group leading to a false inference in an increase in the matrix rank. |
| **qpWave configuration** | **Reliability and Expectation** |
| *Where L = left, R = right* | |
| (L) = L.PP, (R) = R.PP<br>(L) = L.PP, (R) = R.notPP<br>(L) = L.notPP, (R) = R.PP | Reliable. |
| (L) = L.PP, (R) = R.sometimesPP<br>(L) = L.sometimesPP, (R) = R.PP<br>(L) = L .sometimesPP, (R) = R | Unreliable, bias in the matrix of $f_4$ statistics between L.PP and R.PP underlying qpWave could lead to a false estimation of the matrix rank. |

| |
|---|
| .sometimesPP |

**Table S5:** List of all strategies tested to reduce Prime Plus assay bias and their predicted impact on bias. Filters tried with different thresholds have variable values listed inside braces {}.

| **Strategy** | **Hypothesised impact or reason for testing.** |
|---|---|
| Remove transition SNPs | Remove possible impact of ancient DNA damage. |
| Remove reads <50 bp | Reduce impact of reference bias. |
| *Allele frequency-based filters* | |
| SNP missingness {>0%, >50%} | Reduce impact of bias caused by preferential capture of certain SNPs by each enrichment, causing differential patterns of data missingness. |
| Minor allele frequency (MAF) {> 1%, >10%, >30%} | Explore the effect of MAF filters on bias. Informative about the relationship of bias to segregating sites. |
| Linkage disequilibrium filters Window size = 3 variants, slide = 1 variant, $r^2$ = {0.1,0.3,0.5,0.7,0.9} | Remove artefactual 'linkage' between neighbouring variants that could potentially be captured by one bait. Exploratory tested with a range of $r^2$ values due to unknown expected correlation. |
| *SNP-set based filters* | |
| Rohland et al. filter | Reported in Rohland et al. 2022 to allow co-analysis of Legacy 1240k and shotgun data. |
| Filter to Human Origins SNPs | Explorative, to understand the bias impact on these most commonly used SNPs |
| PCA-derived filter | Remove the bias conferred by SNPs driving the "assay bias". |
| *Alternative genotyping* | |
| Random diploid | Explore the capture of alleles at heterozygous sites. |
| Pseudohaploid by majority (>5X sites) | |
| Off-target sites | Remove bias that could be due to the biochemical binding of bait to target DNA, focussing on off-target by-catch. |
| >100 bp out of range sites | Same as above, plus removing biased by-catch where one bait may cover two target sites. |
| *Laboratory protocol adjustments* | |
| Protocol A Single capture round, less amplification cycles | Overall reduction of PCR cycles reduces amplification of any preferential binding bias. |
| Protocol B Single capture round, less amplification cycles, lower | Same as above, and lower hybridisation temperature decreases specificity of bait binding, thus reducing bias in the allele captured. |

| |
|---|
| hybridisation temperature |

**Table S6:** Count of number of SNPs used in each $f_4$ test in Figure 4 and corresponding Z scores.
